# Intraspecific plant-soil feedbacks vary under field conditions among species in a tropical tree community

**DOI:** 10.1101/2023.10.09.561486

**Authors:** Jenalle L. Eck, Lourdes Hernández Hassan, Liza S. Comita

## Abstract

**Premise:** Soil microbes can influence patterns of diversity in plant communities via plant-soil feedbacks. Intraspecific plant-soil feedbacks occur when plant genotype causes variation in soil microbial composition, resulting in differences in the performance of seedlings growing near their maternal plants versus seedlings growing near non-maternal conspecific plants. How commonly such intraspecific plant-soil feedbacks occur in natural plant communities is unclear, especially under variable field conditions.

**Methods:** We conducted an in situ experiment with four native tree species on Barro Colorado Island (BCI), Panama. Seedlings of each species were transplanted beneath their maternal tree or another conspecific tree in the BCI forest. Mortality and growth were assessed at the end of the wet season (∼4 months post-transplant) and at the end of the experiment (∼7 months post-transplant).

**Results:** Patterns of seedling performance varied among species and were inconsistent over time. Significant effects of field environment were detected for two of the four species: at the end of the wet season, *Virola surinamensis* seedlings had higher survival beneath their maternal tree than other conspecific trees, while the opposite pattern was found in *Ormosia macrocalyx.* However, these differences disappeared by the end of the experiment.

**Conclusions:** Our results suggest that intraspecific plant-soil feedbacks occur inconsistently under field conditions in tropical tree species and may have a limited role in determining seedling performance in tropical tree communities. Future studies are needed to elucidate the environmental and genetic factors that determine the incidence and direction of intraspecific plant-soil feedbacks in plant communities.

## INTRODUCTION

Plant-soil feedbacks (PSFs) are a key ecological mechanism by which soil- and plant-associated microbes drive species composition and ecosystem functioning in plant communities worldwide (reviewed by Bever, 2003; Bonanomi et al., 2005; Kulmatiski et al., 2008; Crawford et al., 2019). Plant-soil feedbacks occur due to an accumulation of host-specific microbes, including both mutualists (e.g., mycorrhizal fungi) and antagonists (e.g., microbial pathogens), in the soil around established plants (Bever, 1994; Klironomos, 2002; Packer & Clay, 2003a; reviewed by Ehrenfeld et al., 2005). This causes soil microbial community composition to vary among co-occurring plant species (Westover et al., 1997; Osanai et al., 2013; Burns et al., 2015; Fitzpatrick et al. 2018). Consequently, the soil microbial communities nearby established plants often have differential effects on the survival and/or growth of conspecific versus heterospecific seedlings (see meta-analyses by Kulmatiski et al., 2008; Crawford et al., 2019). However, the potential role of plant genotype in generating PSFs within species is unclear.

Recent experimental evidence suggests that intraspecific PSFs can occur within populations (Eck et al., 2019; Crawford & Hawkes, 2020) and among populations (Bukowski & Petermann, 2014; Liu et al., 2015; Wagg et al., 2015; Bukowski et al., 2018; Kirchoff et al., 2019) of the same plant species. Intraspecific PSFs can be characterized by variation in the performance of closely related seedlings (e.g., offspring) versus unrelated seedlings when exposed to the soil microbial communities of conspecific adults within their population. Intraspecific PSFs may occur if microbes are genotype-specific, i.e., if a given microbial species or community has variable effects on the fitness of different genotypes of a given plant species. This variability may be caused by differences among plant genotypes in their level of resistance to pathogenic species (Alexander et al., 1993; Laine, 2004; reviewed by Gururani et al., 2012) or ability to benefit from mutualistic species (Heath & Tiffin, 2007; Salem et al., 2018; Eck et al., 2022; reviewed by Smith & Goodman, 1999) in their environments. Though genotype-specificity in plant-microbial interactions has been confirmed by molecular studies in several crop and model plant species, equivalent studies in wild plants are scarce. Furthermore, evidence is accumulating that microbial community composition can vary among conspecific plants of different genotype, with corresponding effects on plant fitness or productivity (e.g., Schweitzer et al. 2008; reviewed by terHorst & Zee, 2016). Together, these lines of evidence suggest a potential role for intraspecific PSFs in affecting seedling performance in plant communities.

However, how commonly intraspecific plant-soil feedbacks occur within species in plant communities, and whether they are strong enough to influence seedling performance under field environmental conditions, is unclear. Patterns of seedling performance that fit the predictions of negative intraspecific PSFs have been demonstrated in a handful of plant species (Bukowski & Petermann, 2014; Liu et al., 2015; Bukowski et al., 2018; Eck et al., 2019; Kirchoff et al., 2019; Crawford & Hawkes, 2020), but positive intraspecific PSFs can also occur (Bukowski & Petermann, 2014), as well as lack of intraspecific PSFs in some species (Rallo et al., 2023). Furthermore, existing evidence of intraspecific PSFs arises primarily from experiments conducted under controlled conditions (but see Browne & Karubian, 2016; Kirchoff et al., 2019). In a prior shadehouse experiment, we found that seedlings of a tropical tree species (*Virola surinamensis*) had reduced growth in the soil microbial community from beneath their maternal tree relative to the soil microbial community from beneath non-parent conspecific trees in their population (Eck et al., 2019). Because plant associations with microbes and microbial effects on plant performance can differ between controlled experiments and field conditions (Heinze et al., 2016), it is unclear whether the effects we detected in the shadehouse would translate to patterns of seedling survival in the field. In a field observational study with another tropical tree species, seedlings with rarer genotypes were more likely to survive near conspecific adults than seedlings with more common genotypes (Browne & Karubian, 2016), suggesting that intraspecific PSFs do also occur under field conditions in tropical forests.

Despite the importance of PSFs in shaping the composition and diversity of plant communities, it remains unclear how commonly intraspecific PSFs occur or if they are strong enough to be detected under field conditions. To address this knowledge gap, we conducted a field experiment with four tropical tree species on Barro Colorado Island (BCI), Panama. We asked: does seedling performance differ beneath maternal conspecific trees versus beneath other conspecific trees under field conditions? The field setting of our experiment allowed us to assess seedling performance and test for evidence of intraspecific PSFs in an ecologically relevant context. With this study, we aim to better understand how commonly intraspecific PSFs occur within species in a tropical forest community, allowing better predictions regarding the incidence, magnitude, and direction of intraspecific PSFs and aiding our understanding of their consequences for plant diversity.

## MATERIALS AND METHODS

### Study site and species—

Our study focused on four canopy tree species occurring on Barro Colorado Island (BCI), Republic of Panama (9°09’ N, 79°51’ W). BCI is a 15.6 km^2^ moist tropical lowland forest (Croat, 1978) receiving ∼2600 mm of rainfall per year, with a distinct dry season from ∼January to May (Windsor, 1990). The species represent a range of tropical tree families occurring on BCI: 1) *Lacmellea panamensis* (Woodson) Markgf. (Apocynaceae), 2) *Ormosia macrocalyx* Ducke (Fabaceae), 3) *Tetragastris panamensis* (Engl.) Kuntze (Burseraceae), and 4) *Virola surinamensis* (Rol. ex Rottb.) Warb. (Myristicaceae). *Ormosia macrocalyx*, *T. panamensis*, and *V. surinamensis* are native to much of Central America and northern Amazonia, while *L. panamensis* has a more limited distribution spanning Panama, Costa Rica, and Belize (Croat, 1978). On BCI, the species vary in relative abundance: *T. panamensis* and *V. surinamensis* are common, *L. panamensis* is occasional, while *O. macrocalyx* is rare (Croat, 1978). Seedlings of each species are shade-tolerant (Howe, 1990; Gilbert et al., 2006; Myers & Kitajima, 2007; Krause et al., 2012) but vary in drought sensitivity: *L. panamensis* and *T. panamensis* are drought-tolerant, while *V. surinamensis* is drought-sensitive (Kursar et al., 2009; drought tolerance for *O. macrocalyx* is unknown). Seeds of each species are medium- or large-sized (∼1 – 2 cm) and animal-dispersed, with seed production peaking during March in *T. panamensis* and *L. panamensis*, June in *V. surinamensis*, and September – November and April in *O. macrocalyx* (Croat, 1978; Zimmerman et al., 2007; Wright, S. J., personal communication). *Virola surinamensis* and *T. panamensis* are dioecious, while *L. panamensis* and *O. macrocalyx* are hermaphroditic (Croat, 1978). These species were chosen because their seeds were available at the time of collection and germinated in sufficient quantities in the shadehouse.

### Field experiment—

To test whether seedling performance differed beneath maternal conspecific trees compared to beneath non-maternal conspecific trees, we conducted a field experiment in the BCI forest. We collected seeds from beneath the canopy of six fruiting *V. surinamensis*, three fruiting *T. panamensis*, seven fruiting *L. panamensis*, and three fruiting *O. macrocalyx* on BCI during late June – early August 2015. We assigned the tree that a seed was collected beneath as the seed’s putative mother. Seeds from each parental source were surface sterilized (10 % bleach for 1 m, rinse, 70 % ethanol for 30 s, rinse) and air dried. *Ormosia macrocalyx* seeds were scarified and submerged in water for 24 h prior to planting to encourage germination (Sautu et al., 2007). All seeds were germinated in a shadehouse in autoclaved BCI soil (collected from the forest edge near the shadehouse) under two layers of 80 % shadecloth. One-month post-germination, 145 *V. surinamensis* seedlings, 130 *O. macrocalyx* seedlings, 68 *L. panamensis* seedlings, and 30 *T. panamensis* seedlings were selected for inclusion in the experiment (for a total of 373 experimental seedlings). The number of seedlings per species and seed source reflected availability of seeds collected and healthy seedlings available at the time of transplantation (see Table 1 for an overview of the experimental design).

**Table 1:**
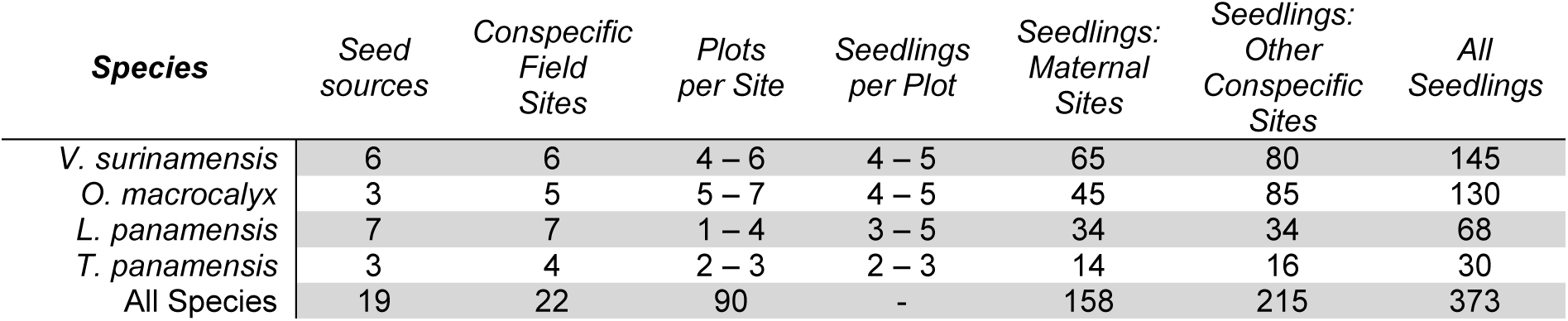
Overview of the field experimental design. In a field experiment on Barro Colorado Island, Panama, we grew seedlings of four tropical tree species (*Virola surinamensis*, *Ormosia macrocalyx*, *Lacmellea panamensis*, and *Tetragastris panamensis*) near their maternal trees or near other conspecific trees in the field. This table provides an overview of our experimental design, including the number of seed sources and conspecific field sites per species, the range in the number of plots per site and the number of seedlings per plot in each species, as well as the number of seedlings planted in maternal sites, in other conspecific sites, and the total number of seedlings per species. Variation among species reflects variation in seed and/or seedling availability at the beginning of the experiment. The total number of plots in the experiment is reported rather than the range of plots per site for all species combined.

| Species | Seed sources | Conspecific Field Sites | Plots per Site | Seedlings per Plot | Seedlings: Maternal Sites | Seedlings: Other Conspecific Sites | All Seedlings |
| --- | --- | --- | --- | --- | --- | --- | --- |
| *V. surinamensis* | 6 | 6 | 4 – 6 | 4 – 5 | 65 | 80 | 145 |
| *O. macrocalyx* | 3 | 5 | 5 – 7 | 4 – 5 | 45 | 85 | 130 |
| *L. panamensis* | 7 | 7 | 1 – 4 | 3 – 5 | 34 | 34 | 68 |
| *T. panamensis* | 3 | 4 | 2 – 3 | 2 – 3 | 14 | 16 | 30 |
| All Species | 19 | 22 | 90 | - | 158 | 215 | 373 |

Seedlings of each species were randomly assigned to one of two treatments: ‘maternal field environment’ or ‘non-maternal conspecific field environment’. Seedlings in the first group were transplanted beneath their own maternal tree in the field. Seedlings in the second group were randomly assigned to be transplanted beneath one of the other seed source trees of their same species (other than their maternal tree). Thus, the experimental treatment signified the putative relationship between an experimental seedling and the conspecific tree it was transplanted beneath during the experiment. Thus, both putative off-spring and non-offspring conspecific seedlings were transplanted beneath each maternal seed source. In the hermaphroditic species, ‘non-maternal conspecific’ trees cannot be ruled out as potential pollen donors to the seedlings. To limit this possibility, in the dioecious species, only female trees were included in the experiment. To increase the number of non-maternal conspecific field environments in *T. panamensis* and *O. macrocalyx*, one additional fruiting *T. panamensis* and two additional fruiting *O. macrocalyx* were selected as seedling transplant sites. All focal trees in the experiment were located by exploring a ∼ 5.5 km^2^ area of BCI with the aid of a mapped 25 ha plot (provided by Wright, S. J., personal communication). Focal trees were located ∼ 100 m to ∼ 4 km apart (see Fig. 1A for map). All met or exceeded the minimum reproductive diameter for their species (Croat, 1978; Wright et al., 2005).

**Fig. 1:**
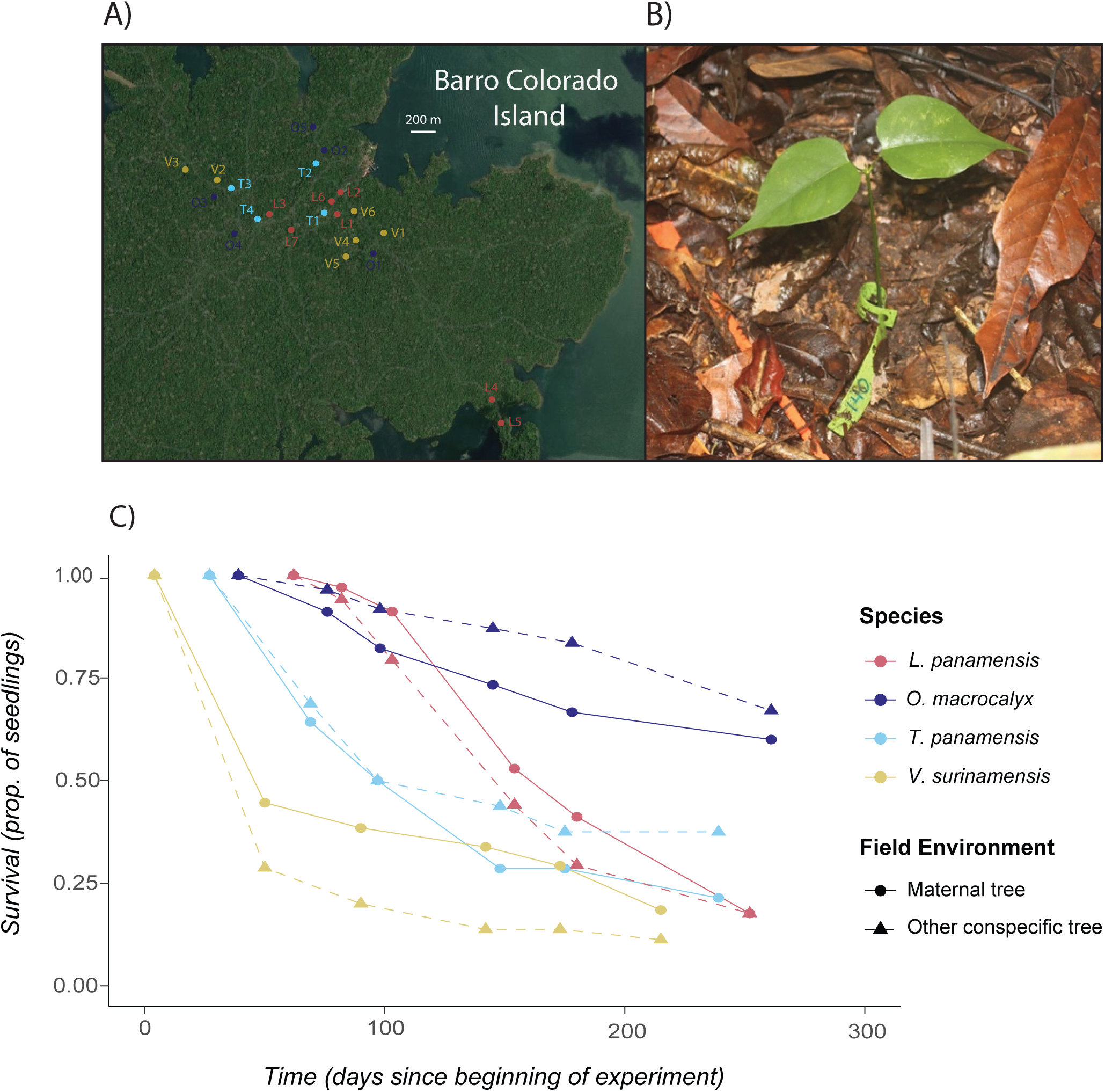
Seedling survival decreased over time in a field experiment on Barro Colorado Island, Panama. *Panel A:* Reproductive trees of four tropical species were used as seed sources and/or seedling transplant sites in a field experiment on Barro Colorado Island (BCI), Panama. Approximate locations of *Virola surinamensis* trees are marked in yellow (V1 – V6), *Lacmellea panamensis* trees in red (L1 – L7), *Ormosia macrocalyx* trees in purple (O1 – O5), and *Tetragastris panamensis* trees in blue (T1 – T4). *Panel B*: In the field experiment, we transplanted seedlings of each species into two experimental treatments: beneath their maternal tree or beneath another conspecific tree in the BCI forest. The photograph shows a stem-tagged *O. macrocalyx* experimental seedling in a field plot on BCI. *Panel C:* The proportion of surviving seedlings declined over time in all species and treatments during the field experiment.

The experiment was set up one species at a time during late August – October 2015. All experimental seedlings were transplanted at ∼ 1 mo of age into their field treatments. Each seedling was randomly transplanted into one of several seedling plots (1 m^2^) beneath the canopy of their assigned conspecific tree. The number of focal plots per tree (1 – 7 plots) and the density of experimental seedlings within each plot was determined by seedling availability: species with more experimental seedlings available had more plots per focal tree and more seedlings per plot (Table 1). Small ranges in the number of conspecific seedlings per plot (2 – 5 seedlings) were used to minimize the potential impact of conspecific seedling neighbors on seedling survival or survival. Plot locations were randomized with respect to direction and distance from the base of the focal tree (1 – 4 m). Each seedling was transplanted into a randomly selected 25-cm^2^ position within a plot. Seedlings roots were not rinsed prior to transplant.

At the time of transplant, each seedling was stem-tagged with a unique identification number (see Fig. 1B for an example photograph). Stem height, the number of leaves, and the length and width of each leaf were also measured for each seedling at the time of transplant. To estimate initial oven-dried biomass for each experimental seedling, we used species-specific allometric equations. These equations arose from models built using measurements of the stem height, leaf area (measured with a leaf area scanner), and total oven-dried biomass of a randomly harvested sample of potential experimental seedlings of each species at the beginning of the experiment (*V. surinamensis*: F_2,47_ = 496.5, p < 0.001, R^2^ = 0.95; *O. macrocalyx*: F_2,37_ = 154.7, p < 0.001, R^2^ = 0.89; *L. panamensis*: F_2,26_ = 183.7, p < 0.001, R^2^ = 0.93; *T. panamensis*: F_2,7_ = 19.7, p = 0.001, R^2^ = 0.81). All field experimental seedlings were censused every 1–2 mo, during which time survival was recorded, and the stem height and number of leaves were measured for all surviving seedlings. We also recorded instances of seedling stem breakage or uprooting (which we refer to as ‘clipping’), likely caused by mammalian herbivores (e.g., agoutis). Seedlings were not watered or enclosed during the field experiment and received only ambient rainfall. Seedlings that were not found during a census (with or without their tags being found) were recorded as dead. After ∼7 mo in the field all surviving seedlings were harvested (*n* = 21 *V. surinamensis* seedlings, *n* = 84 *O. macrocalyx* seedlings, *n* = 12 *L. panamensis* seedlings, and *n* = 9 *T. panamensis* seedlings). Seedlings were harvested one species at a time between late March – early May 2016. Total oven-dried biomass (including aboveground and belowground portions), stem height, and total leaf area (measured with a scanner) were measured at harvest for each surviving seedling. The final months of the experiments overlapped with a severe dry season due to the strong 2015/16 El Niño event: signs of wilting were observed in some seedlings (particularly *V. surinamensis*).

### Statistical analyses—

To test whether seedling performance in the field experiment differed in maternal conspecific field environments relative to non-maternal conspecific field environments, we built a series of mixed-effects models. Because we expected pathogen effects to be stronger in the wet season and because the experiment extended into an unusually severe dry season, we analyzed seedling performance at two time points during the experiment: 1) shortly after the end of the wet season (early-mid January, ∼ 4 months into the experiment), to exclude dry season effects, and 2) at the end of the 7-month experiment (which coincided with the end of the dry season). Seedling performance at each time point was analyzed using data on seedling survival and seedling relative growth rate (based on measurements of seedling stem height and leaf number during the experiment, or biomass at the end of the experiment).

First, we constructed models examining seedling survival at each time point using data from all four species combined. Seedling survival was modelled as a binary response variable using generalized linear mixed-models with binomial errors. ‘Field environment’ (i.e., maternal conspecific or non-maternal conspecific) was included as a fixed effect in each model, as was species. Because we expected that patterns of seedling performance in the field environments could vary among species, we also included an interaction between field environment and species as a fixed effect in these models. Other fixed effects in these models included estimated initial seedling biomass, as well as ‘seedling clipping’ (i.e., whether or not the seedling had experienced clipping up until the time point being analyzed). The identity of the maternal ‘seed source’ tree, the identity of conspecific tree under which the seedling was transplanted (i.e., ‘soil source’), and ‘field plot’ were included as random effects in each model.

To look more deeply at patterns of survival in each species, we then built separate survival models for each of the four species in the experiment during each season. The list of fixed and random effects in these models differed from the model described above in the following ways: 1) ‘species’ and its interaction term with ‘field environment’ were necessarily excluded, 2) due to a limited number of seed sources in *O. macrocalyx* and *T. panamensis*, ‘seed source’ was included in these species’ models as a fixed effect rather than a random effect, and 3) for the same reason, ‘soil source’ was included as a fixed effect rather than a random effect in the *T. panamensis* models. In addition, clipped seedlings and the term denoting clipping were removed from the *O. macrocalyx* and *T. panamensis* models, because clipping was infrequent in these species (4.6 % of seedlings in *O. macrocalyx* and 6.6 % in *T. panamensis*) and including the term prevented convergence in some models. In species where clipping was more frequent, i.e., *V. surinamensis* (31.7 % of seedlings) and *L. panamensis* (16.2 % of seedlings), we built two versions of each model: the first excluded clipped seedlings from the model, while the other retained damaged seedlings but included clipping as a fixed effect. Because including clipped seedlings did not affect model results in *V. surinamensis* and *L. panamensis*, we present the results of the models that include clipped seedlings (i.e., utilizing data on all available seedlings) in these species.

We also tested for differences in seedling relative growth rate (RGR) between the field environments during the experiment. Relative growth rates were calculated based on field measurements of stem height and leaf number of each living seedling during each census interval (Equation 1). Field measurements were converted to estimated seedling biomass (*S*) during each census (*t*) using the species-specific allometric models.

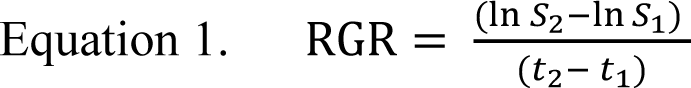

This resulted in a total of 959 RGR observations on 260 seedling individuals across five census intervals. RGR data was analyzed at the end of the wet season and at the end of the experiment using linear mixed-effects models that included data from all four species. The RGR models included the same set of fixed and random effects specified for the survival and biomass models. In addition, the RGR models included the unique identifier of the seedling (to account for repeated measurements of seedlings over time; ‘seedling ID’) and ‘census interval’ as random effects. Because the number of surviving seedlings was small in some species (Fig. 1C), we did not construct separate RGR models for each species. All statistical analyses in our study were conducted in the R statistical environment (R Core Team, 2023). Linear mixed-effects models in our study were constructed using the lme4 package (Bates et al., 2015). P values for mixed-effect model predictors were obtained using the lmerTest package (Kuznetsova et al., 2016) and the car package (Fox & Weisberg, 2019). Long-format data for the RGR analyses was obtained using the tidyr package (Wickham et al., 2023). Plots were generated using the ggplot2 (Wickham, 2016) and interactions (Long, 2019) packages.

## RESULTS

### Seedling survival in the field experiment—

Patterns of seedling survival in maternal field environments versus non-maternal conspecific field environments varied among species at the end of the wet season (Fig. 2A, Table S1), but were similar among species by the end of the experiment (Fig. 2B, Table S1). At the end of the wet season, seedlings of *V. surinamensis* had higher survival when they were growing beneath their maternal tree versus beneath another female conspecific tree (Fig. 3A & Table S2; p = 0.02, *n* = 145 seedlings). In contrast, seedlings of *O. macrocalyx* had lower survival at the end of the wet season when they were growing beneath their maternal tree (Fig. 3C & Table S3; p < 0.05, *n* = 124 seedlings). Differences in seedling survival between field environments in these species disappeared during the dry season (Figs. 3B, 3D, Tables S2, S3). Seedling survival was similar between field environments in *L. panamensis* (Table S4) and *T. panamensis* (Table S5) at both the end of the wet season and end of the experiment. Rates of seedling survival also varied among species (Table S1; *p* < 0.01, *n* = 373 seedlings). Seedlings with higher initial biomass were more likely to survive (Table S1; *p* < 0.01, *n* = 373 seedlings), while clipped seedlings were less likely to survive (Table S1; *p* = 0.02, *n* = 373 seedlings).

**Fig. 2:**
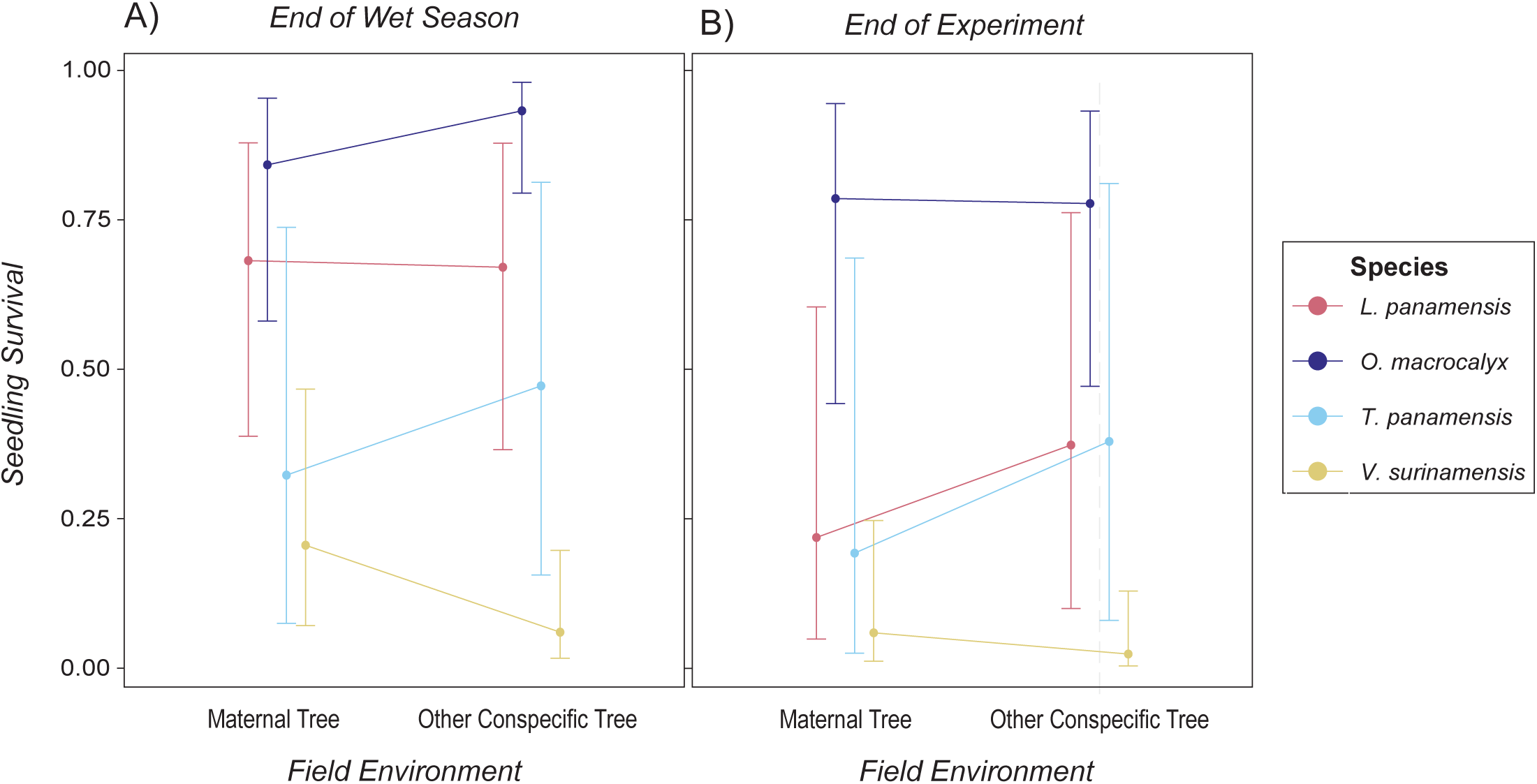
In a field experiment, patterns of seedling survival near maternal conspecific trees and non-maternal conspecific trees varied among species. *Panel A:* In a field experiment on Barro Colorado Island, Panama, field environment (maternal conspecific tree versus non-maternal conspecific tree) and species interacted to jointly determine seedling survival in four focal tropical tree species at the end of the wet season (Table S1; *p* = 0.03 *n* = 373 seedlings). *Panel B*: By the end of the experiment, patterns of seedling survival in the field treatments were similar among species (though species-level differences in survival remained; Table S1; *p* = 0.32, *n* = 373 seedlings). In each panel, the mean estimated survival probability predicted by a generalized linear mixed-effects model is plotted for each species and treatment group. Error bars on the estimates represent a 95 % confidence interval.

**Fig. 3:**
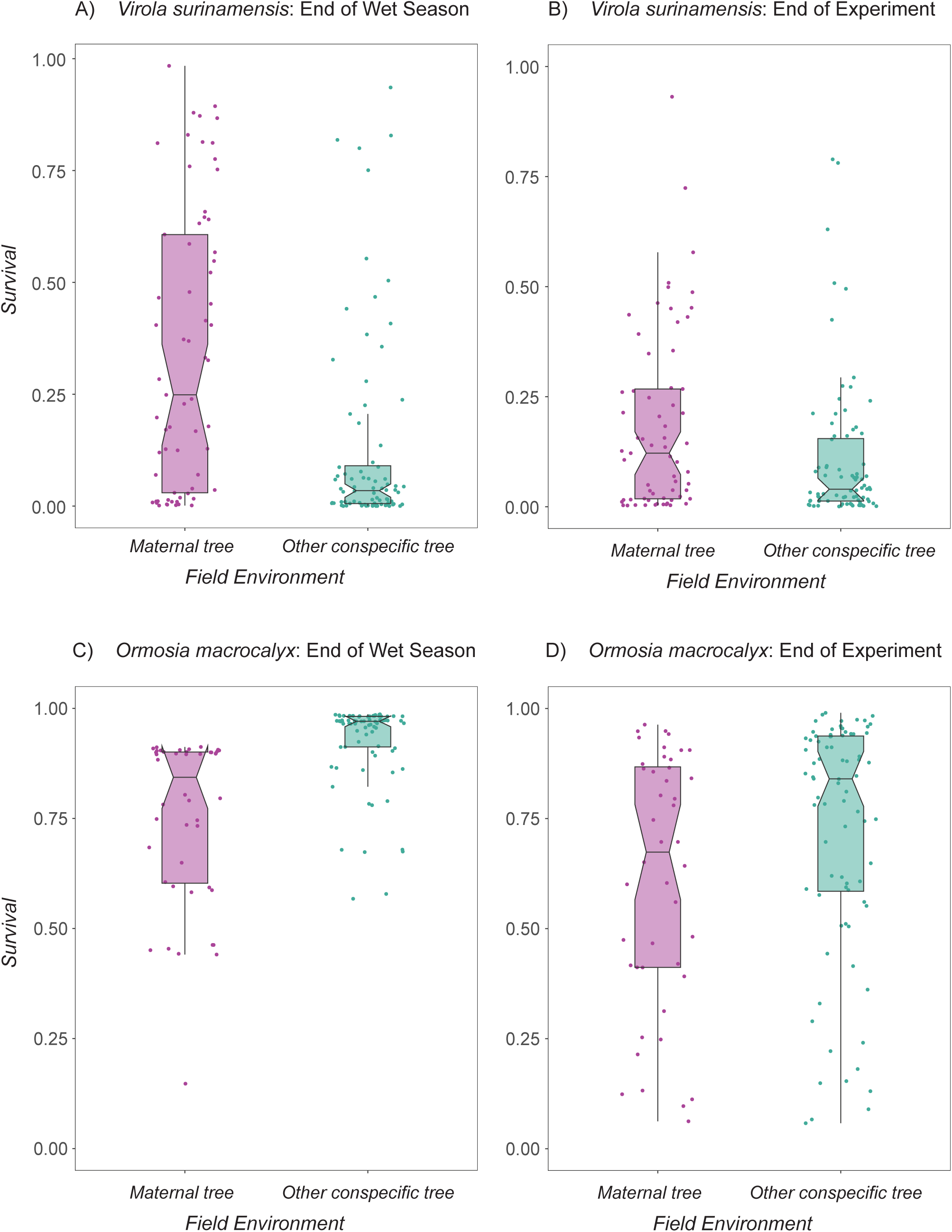
Patterns of seedling survival in *V. surinamensis* and *O. macrocalyx* differed during the field experiment. *Panel A:* In a 7-mo field experiment on Barro Colorado Island, Panama, the survival of *Virola surinamensis* seedlings at the end of the wet season was higher in seedlings beneath their maternal tree than in seedlings beneath a non-parent conspecific female tree (Table S2; *p* = 0.02, *n* = 145 seedlings). *Panel B:* By the end of the experiment, differences in *V. surinamensis* seedling survival were similar among conspecific field environments (Table S2; *p* = 0.25, n = 145 seedlings). *Panel C*: In contrast, survival in *Ormosia macrocalyx* seedlings at the end of the wet season was lower beneath the maternal tree than beneath another conspecific tree (Table S3; *p* = 0.05, *n* = 124 seedlings). *Panel D*: *Ormosia macrocalyx* seedling survival was similar among treatments by the end of the experiment (Table S3; *p* = 0.92, *n* = 124 seedlings). In each panel, dots represent the predicted value for each seedling, box belts show the predicted median values, box notches represent a 95 % confidence interval for comparing predicted medians, box hinges correspond to the first and third quartiles, and box whiskers extend to the largest and smallest value no further than 1.5 × the interquartile range from the hinges.

### Seedling growth in the field experiment***—***

In contrast to patterns of seedling survival, patterns of seedling growth were similar in the field environments during the experiment. Seedling relative growth did not vary between field environments and did not change over time throughout the experiment (Figs. 4A, 4B, Table S6). Relative growth rates varied among species throughout the experiment (Table S6; p < 0.01, *n* = 959 observations). In addition, seedlings with higher initial biomass had higher relative growth rates during the experiment (Table S6; *p* < 0.01, *n* = 959 observations), while clipping reduced seedling relative growth rates (Table S6; *p* < 0.01, *n* = 959 observations).

**Fig. 4:**
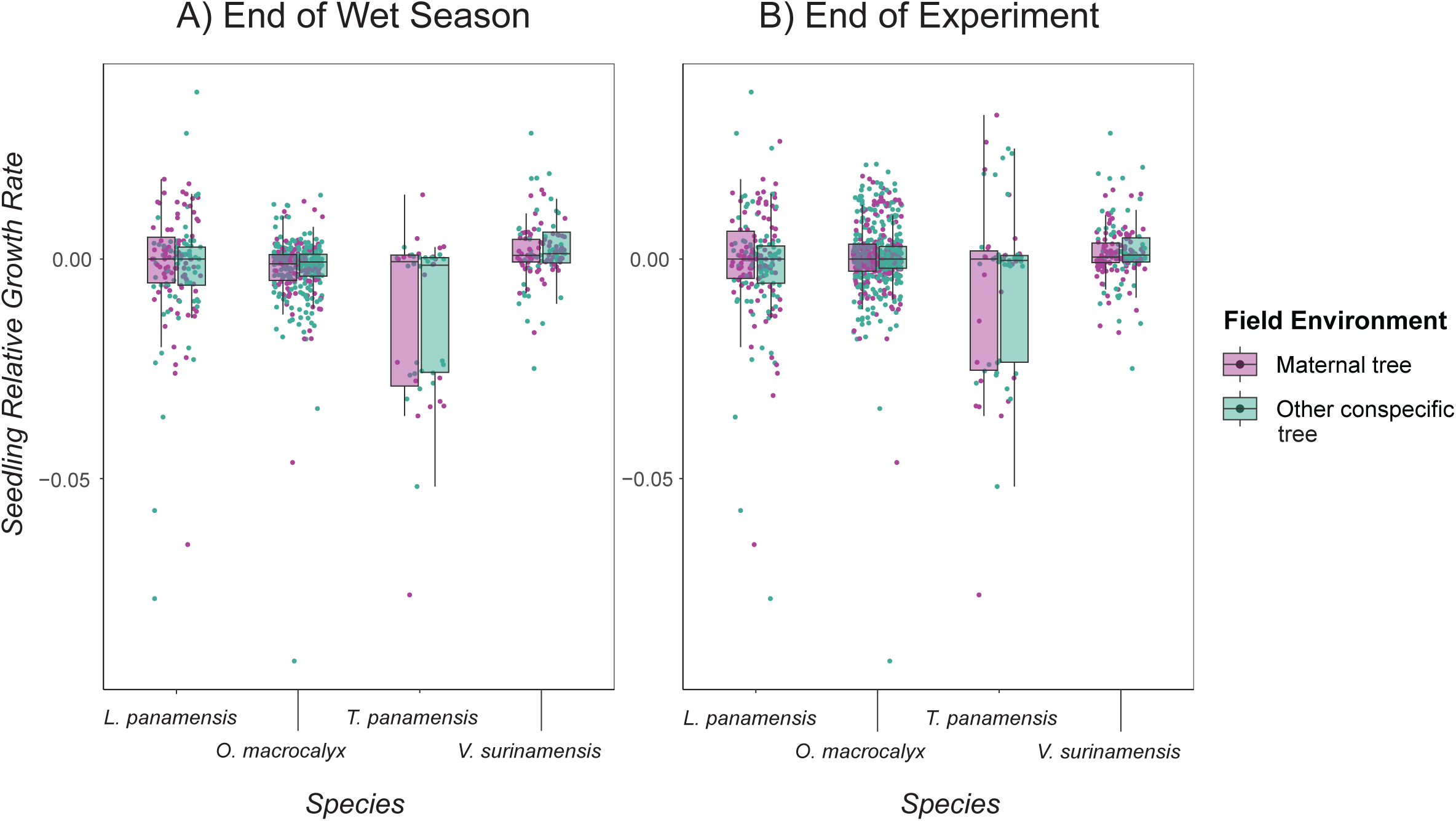
Seedling relative growth rate varied among species but was similar among field environments during the experiment. In a field experiment on Barro Colorado Island, Panama, the relative growth rates of seedlings of four tropical tree species were similar among field environments (maternal tree versus another conspecific tree) at the end of the wet season (*Panel A*; Table S6; *p* = 0.87, n = 668 observations) and at the end of the experiment (*Panel B*; Table S6; *p* = 0.55, *n* = 959 observations). In each panel, dots represent the predicted value for each seedling, box belts show the predicted median values, box hinges correspond to the first and third quartiles, and box whiskers extend to the largest and smallest value no further than 1.5 × the interquartile range from the hinges.

## DISCUSSION

In our in situ field experiment on four tropical tree species, we found that intraspecific plant-soil feedbacks can occur in tropical forests but are inconsistent among species and in time. Though interspecific PSFs are well-documented (e.g., Crawford et al., 2019), relatively few studies have tested for intraspecific PSFs, especially under ecologically-relevant field conditions. In our field experiment in Panama, we detected patterns of seedling performance indicative of intraspecific PSFs in two out of the four focal species. However, the direction of these effects differed between the two species. At the end of the wet season, seedling survival was higher in maternal conspecific field environments relative to non-maternal conspecific field environments in *V. surinamensis* (indicating positive intraspecific PSF), but lower in *O. macrocalyx* (indicating negative intraspecific PSFs). In both species, these patterns were no longer present by the end of the experiment. Thus, our field experiment suggests that intraspecific PSFs can occur, but may play a limited role in determining seedling performance in tropical tree communities.

In a prior shadehouse experiment with the same population of *V. surinamensis* on BCI, we found that seedling growth was reduced in the soil microbial community from beneath their maternal trees relative to in the soil microbial community from beneath other female conspecific trees (Eck et al., 2019). In other words, we found negative intraspecific PSF for growth in the shadehouse, but positive intraspecific PSF for survival in the field in this species. These conflicting findings may reflect differences in conditions in the shadehouse versus field environments. However, they could also indicate that seedlings of this species incur a growth-defense trade off in the soil microbial communities near their maternal tree.

This could occur, for example, if the accumulation of genotype-specific pathogens in the soil near adult trees activates seedling defenses, reducing the resources available for seedling growth but increasing the seedlings’ chances of survival. In the same shadehouse experiment, rates of colonization by arbuscular mycorrhizal fungi were lower in seedlings grown in the soil microbial community of their maternal tree (Eck et al., 2019). As such, dispersed seedlings that grow near another conspecific might avoid genotype-specific pathogens while benefitting from species-specific mycorrhizal fungal growth benefits. The patterns of seedling growth, survival, and mycorrhizal colonization observed in this species could have been produced by an accumulation of genotype-specific pathogens in maternal soils, lack of seedling colonization by mycorrhizal fungi in maternal soils, or by some combination of pathogen and mutualist effects. However, it is important to note that findings from the shadehouse and field experiments might not be directly comparable (Beckman et al., 2022). Additional studies that characterize soil microbial community composition and quantify seedling responses to pathogens and mutualists in the field are needed to better understand these dynamics.

In our study, the four focal tree species varied in patterns of seedling performance in maternal conspecific vs. non-maternal conspecific field environments in the BCI forest. Variation in the incidence, direction, and magnitude of intraspecific PSFs is expected due to variation among plant species in their genetic resources, growth and defensive traits, and life history strategies, alongside microbial differences in host range and effects on plant fitness. Similarly, variation in the incidence, direction, and magnitude of interspecific plant-soil feedbacks among species within plant communities has often been documented (e.g., Mangan et al., 2010; Reinhart, 2012; Bennett et al., 2017). However, characteristics of interspecific PSF might not correlate with characteristics of intraspecific PSF within plant species. In another previous shadehouse experiment with three of our focal species on BCI, *V. surinamensis* and *L. panamensis* exhibited negative interspecific PSFs, while *T. panamensis* did not exhibit interspecific PSF (Mangan et al., 2010). It is difficult to compare studies of interspecific and intraspecific PSF because studies quantifying the former often measure the performance of seedlings of a mixture of genotypes in conspecific soils without distinguishing between parent versus non-parent conspecific sites. It should also be noted that the two species that showed no significant effects of field environment in our study had smaller sample sizes compared to the two species that did show significant effects, so we had limited power to detect significant effects in those species. Nonetheless, we found that seedling performance can differ among conspecific soils in at least some species, and such differences could depend on the level of relatedness between conspecific plants. Genetic relatedness between seedlings and conspecific adult plants has been shown to predict seedling survival in the soil near those adults: soil microbial communities promoted the survival of seedlings from more genetically distant populations (Liu et al., 2015). Variation in seedling performance in conspecific field soils has also been demonstrated to favor the survival of seedlings of locally rare genotypes (Browne & Karubian, 2016) and promote genetic diversity within plant populations (Browne & Karubian, 2018). Together, these studies suggest a role for plant genotype in determining patterns of seedling performance and diversity in the field.

Our findings also indicate that patterns of intraspecific plant-soil feedback could vary over time, potentially due to several factors. First, changing environmental conditions in the field could be responsible for changes in intraspecific PSF over time. In our experiment, patterns of intraspecific PSF emerged during the wet season, but then disappeared by the end of the following dry season. Our experiment coincided with the 2015/2016 El Niño event that resulted in a severe dry season, which was linked to elevated seedling mortality in our study region (Browne et al., 2021). Whether the intraspecific PSFs observed in the wet season persist into the dry season in years without severe droughts is not clear. More generally, environmental variables, such as soil conditions (Wei et al., 2018), plant litter (Veen et al., 2019), and drought (Kaisermann et al., 2017; Fry et al., 2018; reviewed by Hassan et al., 2021; de Vries et al., 2023) have been shown to play a role in determining PSFs (reviewed by van der Putten et al., 2016; De Long et al., 2018). In addition, PSFs are thought to vary temporally (Packer & Clay, 2003b; Hawkes et al., 2013; reviewed by Kardol et al., 2013) and to occur more often in stable relative to variable environments (Duell et al., 2019). Thus, drought and/or seasonality could have affected seedling performance and our end-of-experiment findings. Second, seedling developmental age could have played a role in the patterns we observed. During the experiment, our experimental seedlings aged from approximately one month old to approximately seven months old (though this varied slightly among species). It is possible that intraspecific PSF effects are strongest during certain seedling developmental phases (reviewed by Kardol et al., 2013). Our study would miss any effects of genotype-specific microbes that might impact seed survival, germination, very early seedling mortality, or later-stage seedling performance. Thus, it is possible that other patterns of intraspecific PSF could be present in the species over longer time periods. Though it is challenging to discern the cause of temporal variation in PSFs, the fact that intraspecific PSFs weakened near the end of the experiment in both species that demonstrated them could indicate that this fluctuation was primarily environmental (e.g., potentially caused by overlap with severe dry-season drought conditions) rather than due to changes in the seedlings due to developmental age (which could be expected to vary among species). In addition, the cause of mortality was not usually known for the seedlings in our study, and environmental factors other than soil microbes could have affected seedling survival in our study (Johnson et al., 2017). Additional studies are needed to disentangle the effects of environmental variation versus developmental age on patterns of intraspecific PSFs.

Plant-soil feedbacks allow soil microbes to drive patterns of species diversity in plant communities, but how commonly plant-soil feedbacks occur among genotypes within plant species is unclear, especially under variable field conditions. Intraspecific variability in biotic interactions is often overlooked in community ecology but can greatly affect ecological patterns (reviewed by Freckleton & Lewis, 2006; Bolnick et al., 2011). To date, few studies have tested for evidence of intraspecific plant-soil feedbacks in the field, where seedlings are exposed to their full set of natural enemies (see also Browne & Karubian, 2016; Kirchhoff et al., 2019). Our study highlights the need for molecular studies that characterize soil microbial community composition among plants of known genotype and link this, as well as variation in relatedness, to seedling performance in conspecific sites in the field. Intraspecific variation in seedling performance near conspecific adults could be consequential because of its potential to influence species and/or genetic diversity in plant communities (Stump & Chesson, 2015; Browne & Karubian, 2018; Eck et al., 2019). Future studies are needed to elucidate the environmental, genetic, or trait-based factors that determine the incidence and direction of intraspecific plant-soil feedbacks in plant communities.

## CONCLUSIONS

Understanding the causes and consequences of variability in plant-microbe interactions and plant-soil feedbacks is a key challenge in plant biology. We demonstrate via a field experiment in Panama that the patterns of seedling performance indicative of intraspecific plant-soil feedbacks can occur among species in a tropical forest in Panama but are inconsistent and may ultimately be too weak to detect among other factors that influence seedling survival. Thus, our study suggests a potential, but limited, role of intraspecific plant-soil feedback in determining patterns of seedling performance in tropical tree communities.

## Acknowledgements

The authors thank S. Joseph Wright, E. Allen Herre, E. G. Leigh, Jr., O. Acevedo, B. Jiménez, and S. Murphy for assistance conducting the study in Panama. Funding was provided by National Science Foundation Division of Environmental Biology Grants 1457571 and 1457515 to LSC and European Union ERC Grant 101044424 (PlantSoilAdapt) to Marina Semchenko. JLE acknowledges additional support from Yale University, The Ohio State University, and a Smithsonian Institute Predoctoral Fellowship.

## Author Contributions

JLE and LSC designed the experiments. JLE and LHH set up the experiments and collected the data. JLE analyzed the data with help from LSC. JLE wrote the first draft of the manuscript. All authors contributed to and approved the final version of the manuscript.

## Data Availability Statement

Data are available from the Dryad Digital Repository: https://doi.org/xxx (Eck et al. 202X).

## Literature Cited

1. Alexander, H. M., J. Antonovics, and A. W. Kelly. 1993. Genotypic variation in plant disease resistance – physiological resistance in relation to field disease transmission. Journal of Ecology 81(2): 325–333.

2. Bates, D., M. Maechler, B. Bolker, and W. Walker. 2015. Fitting linear mixed-effects models using lme4. Journal of Statistical Software 67(1): 1–48.

3. Beckman, N. G., R. Dybzinski, and D. Tilman. 2022. Short-term plant-soil feedback experiment fails to predict outcome of competition observed in long-term field experiment. Ecology 104: e3883.

4. Bennett, J. A., H. Maherali, K. O. Reinhart, Y. Lekberg, M. M. Hart, and J. Klironomos. 2017. Plant-soil feedbacks and mycorrhizal type influence temperate forest population dynamics. Science 355: 181–184.

5. Bever, J. D. 1994. Feedback between plants and their soil communities in an old field community. Ecology 75(7): 1965–1977.

6. Bever, J. D. 2003. Soil community feedback and the coexistence of competitors: conceptual frameworks and empirical tests. New Phytologist 157(3): 465–473.

7. Bolnick, D. I., P. Amarasekare, M. S. Araújo, R. Bürger, J. M. Levine, M. Novak, V. H. W. Rudolf, et al. 2011. Why intraspecific trait variation matters in community ecology. Trends in Ecology and Evolution 26(4): 183–192.

8. Bonanomi, G., F. Giannino, and S. Mazzoleni. 2005. Negative plant-soil feedback and species coexistence. Oikos 111: 311–321.

9. Browne, L., and J. Karubian. 2016. Frequency-dependent selection for rare genotypes promotes genetic diversity of a tropical palm. Ecology Letters 19: 1439–1447.

10. Browne, L., and J. Karubian. 2018. Rare genotype advantage promotes survival and genetic diversity of a tropical palm. New Phytologist: doi: 10.1111/nph.15107

11. Browne, L., L. Markesteijn, B. M. J. Engelbrecht, F. A. Jones, O. T. Lewis, E. Manzané-Pinzón, S. J. Wright, and L. S. Comita. 2021. Increased mortality of tropical tree seedlings during the extreme 2015–2016 El Niño. Global Change Biology 27: 5043– 5053.

12. Bukowski, A. R., and J. S. Petermann. 2014. Intraspecific plant-soil feedback and intraspecific overyielding in *Arabidopsis thaliana*. Ecology and Evolution 4(12): 2533–2545.

13. Bukowski, A. R., C. Schittko, and J. S. Petermann. 2018. The strength of negative plant-soil feedback increases from the intraspecific to the interspecific and the functional group level. Ecology and Evolution 8: 2280–2289.

14. Burns, J. H., B. L. Anacker, S. Y. Strauss, and D. J. Burke. 2015. Soil microbial community variation correlates most strongly with plant species identity, followed by soil chemistry, spatial location and plant genus. AoB Plants 7: plv030; doi: 10.1093/aobpla/plv030.

15. Crawford, K. M., and C. V. Hawkes. 2020. Soil precipitation legacies influence intraspecific plant-soil feedback. Ecology 101(10): e03142.

16. Crawford, K. M., J. T. Bauer, L. S. Comita, M. B. Eppinga, D. J. Johnson, S. A. Mangan, S. A. Queenborough, et al. 2019. When and where plant-soil feedback may promote plant coexistence: a meta-analysis. Ecology Letters 22: 1274–1284.

17. Croat, T. B. 1978. Flora of Barro Colorado Island. Stanford University Press, Stanford, CA, USA.

18. De Long, J. R., E. L. Fry, G. F. Veen, and P. Kardol. 2018. Why are plant-soil feedbacks so unpredictable, and what to do about it? Functional Ecology 33: 118–128.

19. Duell, E. B., K. Zaiger, J. D. Bever, and G. W. T. Wilson. 2019. Climate affects plant-soil feedback of native and invasive grasses: negative feedbacks in stable but not in variable environments. Frontiers in Ecology and Evolution 7: 419. doi: 10.3389/fevo.2019.00419.

20. Eck, J. L., S. M. Stump, C. S. Delavaux, S. A. Mangan, and L. S. Comita. 2019. Evidence of within-species specialization by soil microbes and the implications for plant community diversity. PNAS 116(15): 7371–7376.

21. Eck, J. L., M.-M. Kytöviita, and A.-L. Laine. 2022. Arbuscular mycorrhizal fungi influence host infection during epidemics in a wild plant pathosystem. New Phytologist 236: 1922–1935.

22. Eck, J. L., L. Hernández Hassan, and L. S. Comita. 202X. Data from: Intraspecific plant-soil feedbacks vary under field conditions among species in a tropical tree community. – Dryad Digital Repository, xxx.

23. Ehrenfeld, J. G., B. Ravit, and K. Elgersma. 2005. Feedback in the plant-soil system. Annual Review of Environmental Resources 30: 75–115.

24. Fitzpatrick, C. R., J. Copeland, P. W. Wang, D. S. Guttman, P. M. Kotanen, and M. T. J. Johnson. 2018. Assembly and ecological function of the root microbiome across angiosperm plant species. PNAS e1157–e1165. doi: 10.1073/pnas.1717617115

25. Fox, J., and S. Weisberg. 2019. An R Companion to Applied Regression, Third edition. Sage, Thousand Oaks, CA. https://socialsciences.mcmaster.ca/jfox/Books/Companion/.

26. Freckleton, R. P., and O. T. Lewis. 2006. Pathogens, density dependence and the coexistence of tropical trees. Proceedings: Biological Sciences 273(1604): 2909–2916.

27. Fry, E. L., G. N. Johnson, A. L. Hall, W. J. Pritchard, J. M. Bullock, and R. D. Bardgett. 2018. Drought neutralizes plant-soil feedback of two mesic grassland forbs. Oecologia 186: 1113–1125.

28. Gilbert, B., S. J. Wright, H. C. Muller-Landau, K. Kitajima, and A. Hernandéz. 2006. Life history trade-offs in tropical trees and lianas. Ecology 87(5): 1281–1288.

29. Gururani, M. A., J. Venkatesh, C. P. Upadhyaya, A. Nookaraju, S. K. Pandey, and S. W. Park. 2012. Plant disease resistance genes: current status and future directions. Physiological and Molecular Plant Pathology 78: 51–65.

30. Harms, K. E., Condit, R., Hubbell, S. P. and Foster, R. B. 2001. Habitat associations of trees and shrubs in a 50-ha neotropical forest plot. – Journal of Ecology 89: 947–959.

31. Hassan, K., K. M. G. Dastogeer, Y. Carrillo, and U. N. Nielsen. 2021. Climate change-driven shifts in plant-soil feedbacks: a meta-analysis. Ecological Processes 11: 64. doi: 10.1186/s13717-022-00410-z

32. Hawkes, C. V., S. N. Kivlin, J. Du, and V. T. Eviner. 2013. The temporal development and additivity of plant-soil feedback in perennial grasses. Plant and Soil 369: 141–150.

33. Heath, K. D., and P. Tiffin. 2007. Context dependence in the coevolution of plant and rhizobial mutualists. Proc. R. Soc. B 274: 1905–1912.

34. Heinze, J., M. Sitte, A. Schindhelm, J. Wright, and J. Joshi. 2016. Plant-soil feedbacks: a comparative study on the relative importance of soil feedbacks in the greenhouse versus the field. Oecologia 181: 559–569.

35. Howe, H. F. 1990. Survival and growth of juvenile *Virola surinamensis* in Panama: effects of herbivory and canopy closure. Journal of Tropical Ecology 6(3): 259 –280.

36. Johnson, D. J., R. Condit, S. P. Hubbell, and L. S. Comita. 2017. Abiotic niche partitioning and negative density dependence drive tree seedling survival in a tropical forest. Proc. R. Soc. B 284: 20172210. doi: 10.1098/rspb.2017.2210

37. Kardol, P., G. B. De Deyn, E. Laliberté, P. Mariotte, and C. V. Hawkes. 2013. Biotic plant-soil feedbacks across temporal scales. Journal of Ecology 101: 309–315.

38. Kaisermann, A., F. T. de Vries, R. I. Griffiths, and R. D. Bardgett. 2017. Legacy effects of drought on plant-soil feedbacks and plant-plant interactions. New Phytologist 215(4): 1413–1424.

39. Kirchoff, L., A. Kirschbaum, J. Joshi, O. Bossdorf, J. F. Scheepens, and J. Heinze. 2019. Plant-soil feedbacks of *Plantago lanceolata* in the field depend on plant origin and herbivory. Frontiers in Ecology and Evolution 7: 422. doi: 10.3389/fevo.2019.00422

40. Klironomos, J. N. 2002. Feedback with soil biota contributes to plant rarity and invasiveness in communities. Nature 417: 67–70.

41. Krause, G. H., K. Winter, S. Matsubara, B. Krause, P. Jahns, A. Virgo, J. Aranda, and M. García. 2012. Photosynthesis, photoprotection, and growth of shade-tolerant tropical tree seedlings under full sunlight. Photosynth. Res. 113: 273–285.

42. Kulmatiski, A., K. H. Beard, J. R. Stevens, and S. M. Cobbold. 2008. Plant-soil feedbacks: a meta-analytical review. Ecology Letters 11: 980–992.

43. Kursar, T. A., B. M. J. Engelbrecht, A. Burke, M. T. Tyree, B. El Omari, and J. P. Giraldo. 2009. Tolerance to low leaf water status of tropical tree seedlings is related to drought performance and distribution. Functional Ecology 23: 93–102.

44. Kuznetsova, A., P. B. Brockhoff, and R. H. B. Christensen. 2016. lmerTest: Tests in linear mixed effects models. R package version 2.0–33. https://CRAN.R-project.org/package=lmerTest

45. Laine, A.-L. 2004. Resistance variation within and among host populations in a plant-pathogen metapopulation: implications for regional pathogen dynamics. Journal of Ecology 92(6): 990–1000.

46. Liu, X., R. S. Etienne, M. Liang, Y. Wang, and S. Yu. 2015. Experimental evidence for an intraspecific Janzen-Connell effect mediated by soil biota. Ecology 96(3): 662–671.

47. Long, J. A. 2019. Interactions: comprehensive, user-friendly toolkit for probing interactions. R package version 1.1.0, https://cran.r-project.org/package=interactions

48. Mangan, S. A., S. A. Schnitzer, E. A. Herre, K. M. L. Mack, M. C. Valencia, E. I. Sanchez, and J. D. Bever. 2010. Negative plant-soil feedback predicts tree-species relative abundance in a tropical forest. Nature 466: 752–755.

49. Myers, J. A., and K. Kitajima. 2007. Carbohydrate storage enhances seedling shade and stress tolerance in a neotropical forest. Journal of Ecology 95: 383–395.

50. Osanai, Y., D. S. Bougoure, H. L. Hayden, and M. J. Hovenden. 2013. Co-occurring grass species differ in their associated microbial community composition in a temperate native grassland. Plant and Soil 368: 419–431.

51. Packer, A., and K. Clay. 2003a. Soil pathogens and *Prunus serotina* seedling and sapling growth near conspecific trees. Ecology 84(1): 108–119.

52. Packer, A., and K. Clay. 2003b. Development of negative feedback during successive growth cycles of black cherry. Proc. R. Soc. Lond. B 274: 317–324.

53. van der Putten, W. H., M. A. Bradford, E. P. Brinkman, T. F. J. van de Voorde, and G. F. Veen. 2016. Where, when and how plant-soil feedback matters in a changing world. Functional Ecology 30(7): 1109–1121.

54. R Core Team. 2023. R: A language and environment for statistical computing. R Foundation for Statistical Computing, Vienna, Austria. https://www.R-project.org/

55. Rallo, P., S. E. Hannula, F. C. ten Hooven, K. J. F. Verhoeven, J. Kammetenga, and W. H. van der Putten. 2023. Inter- and intraspecific plant-soil feedbacks of grass species. Plant Soil doi: 10.1007/s11104-023-05893-z

56. Reinhart, K. O. 2012. The organization of plant communities: negative plant-soil feedbacks and semiarid grasslands. Ecology 93(11): 2237–2385.

57. Salem, G., M. E. Stromberger, P. F. Byrne, D. K. Manter, W. El-Feki, and T. L. Weir. 2018. Genotype-specific response of winter wheat (*Triticum aestivum* L.) to irrigation and inoculation with ACC deaminase bacteria. Rhizosphere 8: 1–7.

58. Sautu, A., J. M. Baskin, C. C. Baskin, J. Deago, and R. Condit. 2007. Classification and ecological relationship of seed dormancy in a seasonal moist tropical forest, Panama, Central America. Seed Science Research 17: 127–140.

59. Schweitzer, J. A., J. K. Bailey, D. G. Fischer, C. J. LeRoy, E. V. Lonsdorf, T. G. Whitham, and S. C. Hart. 2008. Plant-soil-microorganism interactions: heritable relationship between plant genotype and associated soil microorganisms. Ecology 89(3): 773–781.

60. Smith, K. P., and R. M. Goodman. 1999. Host variation for interactions with beneficial plant-associated microbes. Annual Review of Phytopathology 37: 473–491.

61. Stump, S. M., and P. Chesson. 2015. Distance-responsive predation is not necessary for the Janzen-Connell hypothesis. Theoretical Population Biology 106: 60–70.

62. terHorst, C. P., and P. C. Zee. 2016. Eco-evolutionary dynamics in plant-soil feedbacks. Functional Ecology 30: 1062–1072.

63. Veen, G. F., E. L. Fry, F. C. ten Hooven, P. Kardol, E. Morriën, and J. R. De Long. 2019. The role of plant litter in driving plant-soil feedbacks. Frontiers in Environmental Sciences 7 doi: 10.3389/fenvs.2019.00168.

64. de Vries, F., J. Lau, C. Hawkes, and M. Semchenko. 2023. Plant-soil feedback under drought: does history shape the future? Trends in Ecology and Evolution 38(8): 708–718.

65. Wagg, C., B. Boller, S. Schneider, F. Widmer, and M. G. A. van der Heijden. 2015. Intraspecific and intergenerational differences in plant-soil feedbacks. Oikos 124: 994–1004.

66. Wei, W., M. Yang, Y. Liu, H. Huang, C. Ye, J. Zheng, C. Guo, et al. 2018. Fertilizer N application rate impacts plant-soil feedback in a sanqi production system. Science of the Total Environment 633: 796–807.

67. Westover, K. M., A. C. Kennedy, and S. E. Kelley. 1997. Patterns of rhizosphere microbial community structure associated with co-occurring plant species. Journal of Ecology 85(6): 868–873.

68. Wickham, H. 2016. Ggplot2: elegant graphics for data analysis. Springer-Verlag New York. ISBN 978-3-319-24277-4, https://ggplot2.tidyverse.org

69. Wickham, H., D. Vaughan, and M. Girlich. 2023. Tidyr: tidy messy data. https://tidyr.tidyverse.org.

70. Windsor, D. M. 1990. Climate and moisture variability in a tropical forest: long-term records from Barro Colorado Island, Panamá. – Smithsonian Contributions to the Earth Sciences 29: 1 –145.

71. Wright, S. J., H. C. Muller-Landau, O. Calderón, and A. Hernandéz. 2005. Annual and spatial variation in seedfall and seedling recruitment in a Neotropical forest. Ecology 86(4): 848–860.

72. Zimmerman, J. K., S. J. Wright, O. Calderón, M. A. Pagan, and S. Paton. 2007. Flowering and fruiting phenologies of seasonal and aseasonal neotropical forests: the role of annual changes in irradiance. Journal of Tropical Ecology 23: 231–251.

