## Supplementary Information for "Intraspecific plant-soil feedbacks vary under field conditions among species in a tropical tree community"

The following Supporting Information is available for this article:

**Table S1**. Effect of maternal conspecific versus other conspecific field environments on seedling survival in four species.

**Table S2.** Effect of maternal conspecific versus non-parent female conspecific field environments on *Virola surinamensis* seedling survival.

**Table S3.** Effect of maternal conspecific versus other conspecific field environments on *Ormosia macrocalyx* seedling survival.

**Table S4.** Effect of maternal conspecific versus other conspecific field environments on *Lacmellea panamensis* seedling survival.

**Table S5:** Effect of maternal conspecific versus non-parent female conspecific field environments on *Tetragastris panamensis* seedling survival.

**Table S6.** Effect of maternal conspecific versus other conspecific field environments on seedling relative growth rates in four species.

**Table S1: Effect of maternal conspecific versus other conspecific field environments on seedling survival in four species.** We tested whether the survival of seedlings of four species varied among field environments (maternal conspecific tree versus other conspecific tree) during a 7-mo field experiment on Barro Colorado Island using two generalized linear mixed-effects models ($n$ = 373 seedlings). In the first model, seedling survival was analyzed at the end of the wet season. In the second model, seedling survival was analyzed at the end of the experiment. Type II analysis of deviance results from Wald chi-square tests are shown below, including chi-squared (*Chisq*), degrees of freedom (*d.f.*) and p-values (*p*). Fixed effects for which $p \leq 0.05$ are in bold. The variance and standard deviation for each random effect in the models is also reported.

| Analysis of Deviance | | End of Wet Season | | End of Experiment | |
|---|---|---|---|---|---|
| ***Fixed Effect*** | *d.f.* | *Chisq* | *p* | *Chisq* | *p* |
| Field Environment | 1 | 0.30 | 0.59 | 0.02 | 0.90 |
| Species | 3 | 21.47 | **8.42 x $e^{-05}$** | 15.73 | **1.29 x $e^{-03}$** |
| Initial Seedling Biomass | 1 | 14.07 | **1.76 x $e^{-04}$** | 18.61 | **1.61 x $e^{-05}$** |
| Seedling Clipping | 1 | 5.27 | **0.02** | 7.80 | **0.01** |
| Field Environment × Species | 3 | 9.30 | **0.03** | 3.49 | 0.32 |

| ***Random Effect*** | End of Wet Season | | End of Experiment | |
|---|---|---|---|---|
| | *Variance* | *Std. Dev.* | *Variance* | *Std. Dev.* |
| Seed Source | 0.02 | 0.13 | 0.01 | 0.07 |
| Soil Source | 0.95 | 0.97 | 1.55 | 1.24 |
| Field Plot | 0.64 | 0.80 | 1.50 | 1.22 |

**Table S2: Effect of maternal conspecific versus non-parent female conspecific field environments on *Virola surinamensis* seedling survival.** We tested whether the survival of *V. surinamensis* seedlings during a 7-mo field experiment on Barro Colorado Island varied among field environments (maternal conspecific tree versus non-parent female conspecific tree) using two generalized linear mixed-effects models ($n$ = 145 seedlings). In the first model, seedling survival was analyzed at the end of the wet season. In the second model, seedling survival was analyzed at the end of the experiment. Type II analysis of deviance results from Wald chi-square tests are shown below, including chi-squared (*Chisq*), degrees of freedom (*d.f.*) and p-values (*p*). Fixed effects for which $p \leq 0.05$ are in bold. The variance and standard deviation for each random effect in the models is also reported.

| **Analysis of Deviance** | | **End of Wet Season** | | **End of Experiment** | |
|---|---|---|---|---|---|
| ***Fixed Effect*** | *d.f.* | *Chisq* | *p* | *Chisq* | *p* |
| Field Environment | 1 | 5.46 | **0.02** | 1.30 | 0.25 |
| Initial Seedling Biomass | 1 | 10.18 | **1.42 x $e^{-03}$** | 9.17 | **2.47 x $e^{-03}$** |
| Seedling Clipping | 1 | 2.14 | 0.14 | 5.10 | **0.02** |

| ***Random Effect*** | **End of Wet Season** | | **End of Experiment** | |
|---|---|---|---|---|
| | *Variance* | *Std. Dev.* | *Variance* | *Std. Dev.* |
| Seed Source | 0.85 | 0.92 | 0.00 | 0.00 |
| Soil Source | 7.45 | 2.73 | 3.73 | 1.93 |
| Field Plot | 1.61 | 1.27 | 0.59 | 0.77 |

**Table S3: Effect of maternal conspecific versus other conspecific field environments on *Ormosia macrocalyx* seedling survival.** We tested whether the survival of *O. macrocalyx* seedlings during a 7-mo field experiment on Barro Colorado Island varied among field environments (maternal conspecific tree versus other conspecific tree) using two generalized linear mixed effects models ($n$ = 124 seedlings). In the first model, seedling survival was analyzed at the end of the wet season. In the second model, seedling survival was analyzed at the end of the experiment. Clipped seedlings were excluded from each model. Type II analysis of deviance results from Wald chi-square tests are shown below, including chi-squared (*Chisq*), degrees of freedom (*d.f.*) and p-values (*p*). Fixed effects for which $p \leq 0.05$ are in bold. The variance and standard deviation for each random effect in the models is also reported.

| **Analysis of Deviance** | | **End of Wet Season** | | **End of Experiment** | |
|---|---|---|---|---|---|
| ***Fixed Effect*** | *d.f.* | *Chisq* | *p* | *Chisq* | *p* |
| Field Environment | 1 | 4.00 | **0.05** | 0.01 | 0.92 |
| Initial Seedling Biomass | 1 | 0.02 | 0.88 | 7.39 | **0.01** |
| Seed Source | 1 | 0.89 | 0.64 | 3.52 | 0.17 |

| ***Random Effect*** | **End of Wet Season** | | **End of Experiment** | |
|---|---|---|---|---|
| | *Variance* | *Std. Dev.* | *Variance* | *Std. Dev.* |
| Soil Source | 0.68 | 0.83 | 1.27 | 1.13 |
| Field Plot | 1.79 | 1.34 | 3.20 | 1.79 |

**Table S4: Effect of maternal conspecific versus other conspecific field environments on *Lacmellea panamensis* seedling survival.** We tested whether the survival of *L. panamensis* seedlings during a 7-mo field experiment on Barro Colorado Island varied among field environments (maternal conspecific tree versus other conspecific tree) using two generalized linear mixed effects models ($n$ = 68 seedlings). In the first model, seedling survival was analyzed at the end of the wet season. In the second model, seedling survival was analyzed at the end of the experiment. Type II analysis of deviance results from Wald chi-square tests are shown below, including chi-squared (*Chisq*), degrees of freedom (*d.f.*) and p-values (*p*). Fixed effects for which $p \leq 0.05$ are in bold. The variance and standard deviation for each random effect in the models is also reported.

| **Analysis of Deviance** | | **End of Wet Season** | | **End of Experiment** | |
|---|---|---|---|---|---|
| ***Fixed Effect*** | *d.f.* | *Chisq* | *p* | *Chisq* | *p* |
| Field Environment | 1 | 0.18 | 0.67 | 0.03 | 0.85 |
| Initial Seedling Biomass | 1 | 2.04 | 0.15 | 1.36 | 0.24 |
| Seedling Clipping | 1 | 1.08 | 0.30 | 0.23 | 0.63 |

| ***Random Effect*** | **End of Wet Season** | | **End of Experiment** | |
|---|---|---|---|---|
| | *Variance* | *Std. Dev.* | *Variance* | *Std. Dev.* |
| Seed Source | 0.00 | 0.00 | 0.05 | 0.23 |
| Soil Source | 0.59 | 0.77 | 2.11 | 1.45 |
| Field Plot | 0.00 | 0.00 | 0.00 | 0.00 |

**Table S5: Effect of maternal conspecific versus non-parent female conspecific field environments on *Tetragastris panamensis* seedling survival.** We tested whether the survival of *T. panamensis* seedlings during a 7-mo field experiment on Barro Colorado Island varied among field environments (maternal conspecific tree versus non-parent female conspecific tree) using two generalized linear mixed effects models ($n$ = 30 seedlings). In the first model, seedling survival was analyzed at the end of the wet season. In the second model, seedling survival was analyzed at the end of the experiment. Clipped seedlings were excluded from each model. Type II analysis of deviance results from Wald chi-square tests are shown below, including chi-squared (*Chisq*), degrees of freedom (*d.f.*) and p-values (*p*). Fixed effects for which $p \leq 0.05$ are in bold. The variance and standard deviation for each random effect in the models is also reported.

| **Analysis of Deviance** | | **End of Wet Season** | | **End of Experiment** | |
|---|---|---|---|---|---|
| ***Fixed Effect*** | *d.f.* | *Chisq* | *p* | *Chisq* | *p* |
| Field Environment | 1 | 0.28 | 0.60 | 2.14 | 0.14 |
| Initial Seedling Biomass | 1 | 0.49 | 0.48 | 1.20 | 0.27 |
| Seed Source | 2 | 2.93 | 0.23 | 5.39 | 0.07 |
| Soil Source | 3 | 2.79 | 0.43 | 6.59 | 0.09 |

| ***Random Effect*** | **End of Wet Season** | | **End of Experiment** | |
|---|---|---|---|---|
| | *Variance* | *Std. Dev.* | *Variance* | *Std. Dev.* |
| Field Plot | 0.89 | 0.94 | 2309 | 48.05 |

**Table S6: Effect of maternal conspecific versus other conspecific field environments on seedling relative growth rates in four species.** We tested whether the relative growth rates of seedlings of four species varied among field environments (maternal conspecific tree versus non-parent female conspecific tree) during a 7-mo field experiment on Barro Colorado Island using two linear mixed-effects models. In the first model, seedling relative growth rates were analyzed at the end of the wet season ($n$ = 668 observations). In the second model, seedling relative growth rates were analyzed at the end of the experiment ($n$ = 959 observations). Type II analysis of deviance results from Wald chi-square tests are shown below, including chi-squared (*Chisq*), degrees of freedom (*d.f.*) and p-values (*p*). Fixed effects for which p $\leq$ 0.05 are in bold. The variance and standard deviation for each random effect in the models is also reported.

| **Analysis of Deviance** | | **End of Wet Season** | | **End of Experiment** | |
|---|---|---|---|---|---|
| ***Fixed Effect*** | *d.f.* | *Chisq* | *p* | *Chisq* | *p* |
| Field Environment | 1 | 0.03 | 0.87 | 0.35 | 0.55 |
| Species | 3 | 109.37 | **< 2.2 x $e^{-16}$** | 61.71 | **2.53 x $e^{-13}$** |
| Initial Seedling Biomass | 1 | 8.41 | **3.73 x $e^{-03}$** | 8.41 | **3.73 x $e^{-03}$** |
| Seedling Clipping | 1 | 55.66 | **8.63 x $e^{-14}$** | 39.58 | **3.14 x $e^{-10}$** |
| Field Environment × Species | 3 | 1.18 | 0.76 | 1.43 | 0.70 |

| ***Random Effect*** | **End of Wet Season** | | **End of Experiment** | |
|---|---|---|---|---|
| | *Variance* | *Std. Dev.* | *Variance* | *Std. Dev.* |
| Seed Source | 0.00 | 0.00 | 1.55 x $e^{-14}$ | 1.24 x $e^{-07}$ |
| Soil Source | 0.00 | 0.00 | 0.00 | 0.00 |
| Field Plot | 0.00 | 0.00 | 0.00 | 0.00 |
| Seedling ID | 0.00 | 0.00 | 0.00 | 0.00 |
| Census Interval | 2.98 x $e^{-06}$ | 1.73 x $e^{-03}$ | 3.20 x $e^{-05}$ | 0.01 |
| Residual | 9.40 x $e^{-05}$ | 0.01 | 8.22 x $e^{-05}$ | 0.01 |
